# Brain-wide representational drift: memory consolidation and entropic force

**DOI:** 10.1101/2025.09.25.678547

**Authors:** Felipe Yaroslav Kalle Kossio, Raoul-Martin Memmesheimer

## Abstract

Memory engrams change on the microscopic level with time and experience as the neurons that compose them switch in a process termed representational drift. On the macroscopic level the engrams are not static either: numbers of engram neurons in different brain regions change over time, which is considered to reflect the process of memory consolidation. Here we predict a link between these two levels, using a novel statistical physics approach to engram modeling. Importantly, it is general as it makes only minimal assumptions on the engram’s nature, does not rely on a specific architecture, and its fundamental implications hold for any representational drift with random component. Our first, analytically well tractable model shows that an entropic force emerges at the region level from random representational drift at the neuronal level. We refine the model by incorporating the interaction of the entropic force with biological processes that shape neuronal engrams. Here, the connectivity between the brain regions strongly influences the (quasi-)equilibrium engram distribution. The obtained distributions are in qualitative agreement with the ones of a biologically detailed drifting assembly model. Our engram description allows to predict the engram evolution in large neuronal systems such as the mouse brain. We find several predictions that are consistent across the valid tested parameter ranges, such as a strong tendency of the engram to leave the hippocampus. The results suggest that the brain operates in a regime where engrams drift, both deterministically and randomly, to allow for memory consolidation.

## Introduction

One of the most important functions of the brain is to convert experiences into memories and to store these memories for significant amounts of time. Ensembles of neurons and the synapses interlinking them are believed to provide the physical substrate for the memory storage: the engram [1–6]. Memory engrams are not static: On the microscopic level the neurons that compose them change with time and experience in a process termed representational drift [3, 7–9], whose functional role is still unclear. On the macroscopic level the numbers of engram neurons in different brain regions change over time [10,11]. Furthermore, over time memories are consolidated [12–14]; consequently, the macroscopic change of engram neuron numbers is often assumed to be a neuronal correlate of memory consolidation.

A classical view of consolidation posits that the parts of a new memory engram, which are distributed throughout the brain, are crucially connected through the engram parts in the hippocampus [13, 15, 16]. The hippocampus then guides the isocortex to form direct or indirect connections between the parts itself. Over time, the memory may thereby become completely independent of the hippocampus. This view of consolidation inspired a large number of modeling studies [17–26]. These highlight a variety of network architectures and plasticity rules that may allow the transfer of the functionality of the hippocampal engram parts to the isocortex. However, there are a number of weaknesses of the classical view of consolidation and alternative possibilities exist [27].

Here we develop models connecting representational drift and the distribution of an engram across the brain’s regions over the course of time. In our models the memory engrams drift by deterministic and random remodeling. Importantly, the random remodeling can induce on a macroscopic scale practically deterministic transitions between brain areas. Our models do not require the hippocampus to guide the isocortex during consolidation. The hippocampus might then just be one region among many others, which initially binds distributed engram parts possibly due to its connectivity and specific learning abilities [28]. Using an approach from statistical physics as well as detailed neural network modeling we show how the random drift of a memory engram at the microscopic level leads to the emergence of a practically deterministic entropic force [29] on the macroscopic engram, which tends to equilibrate memory coding levels across the brain regions. We further show how neuronal preferences, which can be biologically implemented via plasticity rules and govern deterministic drift, may be captured by an energy function. This allows to study their interaction with the random drift within our framework. We then introduce a detailed biological model of a drifting engram and find that its evolution characteristics qualitatively agree with those of the statistical physics models. Finally we apply our model of engram drift to the whole mouse brain to model and thereby predict macroscopic engram transformation. Our findings suggest that the brain operates in the regime that supports representational drift to allow memory consolidation.

## Results

### A purely-random engram drift

Engram tracking experiments often report the number of engram neurons in a particular brain region [10, 11]. We call the state specified by this macroscopic level description the macrostate. It omits the exact microscopic details of the engram structure. Consider first a single brain region and an engram within it. The macrostate is then simply the size of the engram: the number *n* of neurons that form it, Fig. 1. A more detailed description of the engram, listing the particular neurons forming it, specifies a microstate. The same engram macrostate can thus result from many different microstates. We imagine each engram neuron to possess strong synaptic connections with the rest of the engram [4, 5, 30], without yet explicitly modeling them.

**Figure 1:**
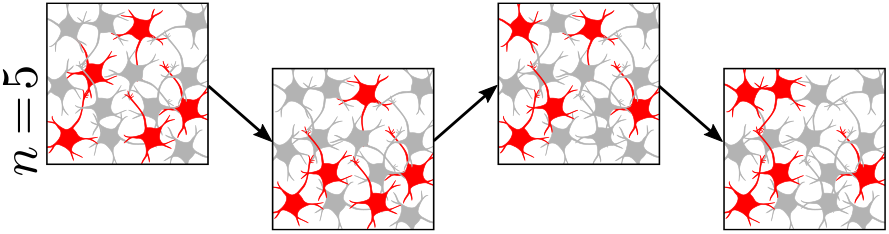
Engram drifting in a single brain region. Engram neurons are shown in red; non-engram neurons in gray. In our first, simple statistical model, the microstate performs a random walk with conserved overall engram size, here *n* = 5. If there is only a single brain region, the engram size defines the macrostate, which is therefore also conserved.

Experiments show that over time, some neurons may leave the engram and others join it, such that the engram drifts [3, 8, 9]. Such representational drift may be caused, for example, by noise in the plasticity rules, spontaneous synaptic turnover, or change of intrinsic properties [31–33]. The exact mechanism behind the drift is not important for our models and analysis. As a first, simple model, we assume that at each time evolution step, a random neuron leaves the engram and a random neuron joins it. Then, the engram size *n* and, in the case of a single region, the macrostate is conserved, Fig. 1. The drift is a random walk through the microstates. Mathematically it is a random walk on a Johnson graph [34, 35]. Since each microstate has the same number of microstates that it can transition to and arise from, each node of the corresponding graph has the same number of connections. This implies that in the long run each microstate has the same frequency of occurrence [34]. Since all microstates are equally likely, in statistical physics terms they form a microcanonical ensemble [36].

### Engram drifting in two brain regions

For two brain regions, an engram has *n*_1_ neurons in the first region and *n*_2_ neurons in the second region, Fig. 2a; *n*_1_ and *n*_2_ define its macrostate ***n*** = (*n*_1_, *n*_2_). Random representational drift may gradually change the ensemble of engram neurons and thereby the microstate ***m*** (*m*_*i*_ = 1 if neuron *i* is an engram neuron and 0 otherwise). However, the macrostate now changes even if the overall engram size stays constant, Fig. 2b: this is because, for example, a neuron in Region 1 may leave the engram and a neuron in Region 2 may join it.

**Figure 2:**
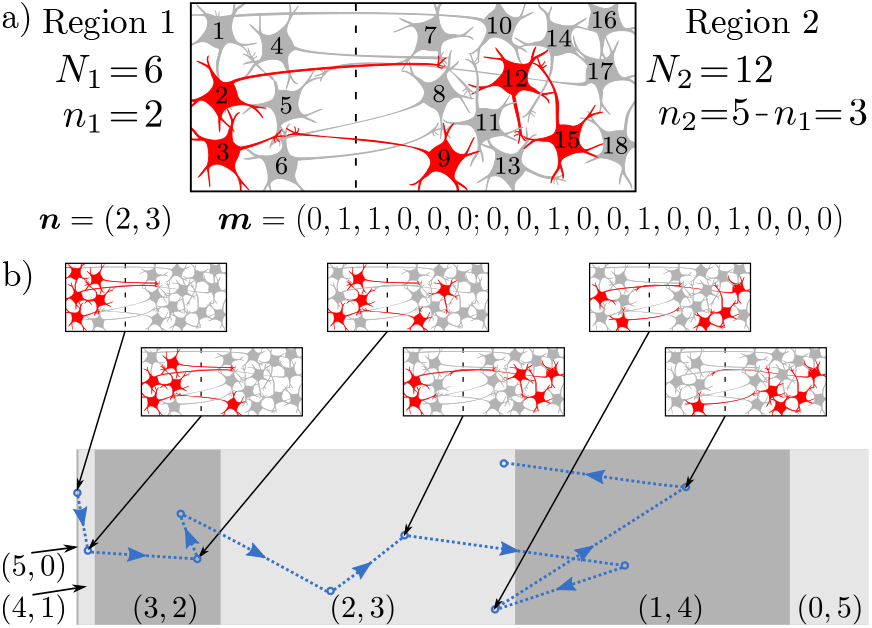
Engram drifting in two brain regions. (a) Example engram microstate for two regions with 6 and 12 neurons (dashed line: separation between the regions). The engram contains five neurons, shown in red; non-engram neurons are displayed in gray. To define a microstate, each neuron is assigned an index. The *i*th entry of the vector ***m*** characterizing the microstate is 1 if the neuron with index *i* is part of the engram and 0 otherwise. The two entries of the vector ***n*** characterizing the macrostate are the numbers of engram neurons in the two regions. (b) Schematic engram trajectory due to drift. (b, upper) Illustration of the realized microstates. (b, lower) Trajectory in the microstate space. Gray shaded areas indicate macrostates and their multiplicities Ω(***n***).

The simplest way to obtain a two-region network is to superficially define two regions in a single region network, without applying any further changes. We chose such a partition into a smaller region containing *N*_1_ neurons and a larger one with *N*_2_ neurons, where *N*_1_ + *N*_2_ = *N* is the total network size. We consider a more biologically motivated partition in a later section. In the context of classical theories of memory consolidation (see Introduction), the smaller region may be interpreted as hippocampus and the larger one as isocortex. Assuming again that at each step a random neuron leaves the engram and a random neuron joins it, the process is a random walk in the microstate space with the same properties as in the case of a single region. In particular, in the long run each accessible microstate occurs with the same frequency. The probability of an engram being in a particular macrostate is then proportional to the “volume” that the macrostate occupies in the microstate space, more precisely to its multiplicity Ω(***n***), i.e. to the number of underlying microstates, Fig. 2b. This is the origin of the entropic force that distributes the engram over different brain regions in our models.

Importantly, the entropic force leads to directed engram changes on the macroscale. To clarify this, we first study the extreme case where engram neurons are initially only in the smaller region; more realistic initial states are considered in a later section. Fig. 2b exemplarily displays engram micro- and macrostates during the initial phase of drift in a very small network. Fig. 3a shows the evolution of the engram macrostates due to drift in a larger network. We observe that the majority of the engram quickly leaves the smaller region (the hippocampus) and enters the larger one (the isocortex) until equilibrium, Fig. 3b, is reached. This general engram transformation may explain the similar experimentally observed transformation that is often identified with memory consolidation and transient initial dependence of many memories on the hippocampus [12–14, 27, 37].

**Figure 3:**
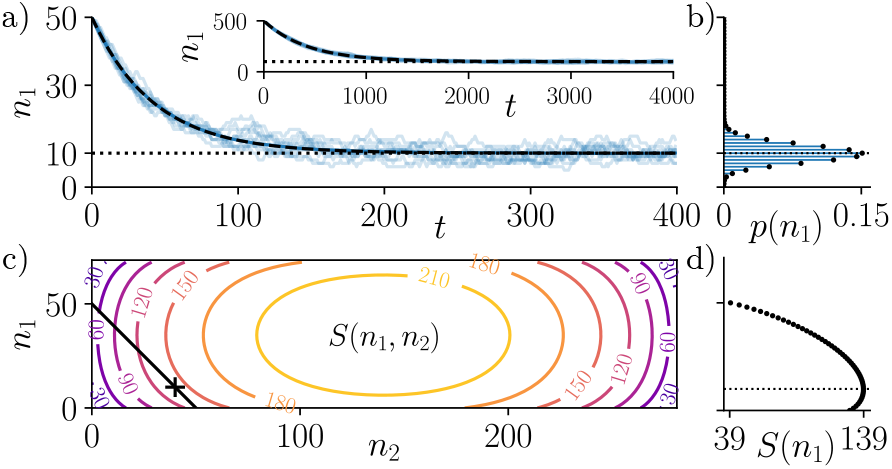
Drift of an engram of fixed size (*n* = 50) in a network with two regions (*N*_1_ = 70, *N*_2_ = 280). (a) Ten example engram drift trajectories (light blue). The engrams are initially completely in Region 1, *n*_1_(*t* = 0) = *n*. They quickly enter Region 2 and eventually settle at equilibrium. The majority of engram neurons is then in Region 2. Individual trajectories stay near the ensemble average (blue) and the matching theoretical average (dashed black); the time evolution is close to deterministic despite the rather small size of the considered engram. Time *t* is the number of engram neuron replacements. (Inset) In a ten times larger system the relative size of fluctuations around the mean trajectory is smaller such that they are hardly visible. (b) Theoretical (black) and observed (blue, 1000 samples) equilibrium probability distributions of a macrostate with *n*_1_ engram neurons in Region 1. (c) Entropy *S*(***n***) of the macrostate ***n*** = (*n*_1_, *n*_2_). The black line indicates the accessible macrostates, which satisfy the constraint *n* = *n*_1_ + *n*_2_; the cross indicates the average of the equilibrium distribution. The microstates are discrete but dense, we use lines instead of points for visualization only. (d) Entropy of the accessible macrostates, *S*(*n*_1_) = *S*(*n*_1_, *n* − *n*_1_), i.e. essentially the entropy along the black line in (b, left). The thin dotted lines in (a,b,d) highlight the equilibrium distribution’s average, which is nearly identical to the most probable macrostate.

### Engram equilibrium

Since in our simple model the engram size is fixed After a sufficiently long period of drift the probability that the engram is in a particular macrostate ***n*** = (*n*_1_, *n*_2_) is proportional to its multiplicity. We can obtain this number of corresponding microstates by computing the number of ways to select *n*_1_ engram neurons from the *N*_1_ available Region 1 neurons and *n*_2_ engram neurons from the *N*_2_ available Region 2 neurons, 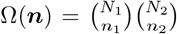 Since in our simple model the engram size is fix to *n*, only those macrostates are accessible (can be present) that satisfy the constraint *n*_1_ + *n*_2_ = *n*. The multiplicity is typically a very large number. Therefore we often take its logarithm, ln Ω(***n***), which can be identified with the Boltzmann entropy from statistical physics, *S*(***n***) = ln Ω(***n***), Fig. 3c,d.

The macrostate equilibrium probability, Figure 3b, is *p*(***n***) = Ω(***n***)*/*Ω(*n*) where the normalizing factor is one over the total number of accessible microstates,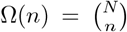. The expected equilibrium number of engram neurons in the first region is ⟨*n*_1_⟩_eq_ = *nN*_1_*/N* . This makes sense intuitively: At a time point long after equilibration, the engram will be homogeneously distributed over the network. Thus the expected fraction of engram neurons in Region 1, ⟨*n*_1_⟩_eq_*/n* equals the fraction of neurons in Region 1, *N*_1_*/N* . The drift thus tends to equalize memory coding levels, i.e. fractions of engram neurons, between the regions: As an example in our case of two regions we get ⟨*n*_1_⟩_eq_*/N*_1_ = ⟨*n*_2_⟩_eq_*/N*_2_.

The fluctuations at equilibrium, characterized by the coefficient of variation, are 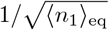 (for large engrams and networks, Methods): As the number of engram neurons in a region becomes larger, the relative size of equilibrium fluctuations decreases and the equilibrium distribution becomes more sharply peaked.

### Approach to the equilibrium

We now examine the evolution of far-from-equilibrium engram macrostates in more detail. Starting in some fixed initial macrostate *n*_1_(0), the engram reaches macrostate *n*_1_(*t*) after *t* time steps. The average macrostate trajectory due to drift is

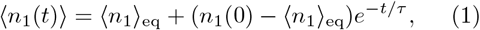

where 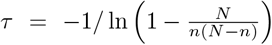 (Methods) and the fluctuations around the average trajectory are small, see Fig. 3a. Therefore, the microscopically random engram drift leads on the macroscopic scale to a directed evolution towards equilibrium. For *N* ≫*n* ≫ 1, which we expect for typical systems, the exponential relaxation to the equilibrium is determined just by the engram size, *τ ≈ n*. The emergence of practically deterministic macroscopic dynamics in a sufficiently large stochastic system is well known in statistical physics: it marks, for example, the transition to thermodynamics [36, 38]. In our system, the directed evolution suggests that there is a force acting on the engram, which drives it towards the equilibrium state. This force is the emergent entropic force [29]. It originates from the fact that the engram remodels randomly and has a large number of possible microstates.

Importantly we demonstrated the existence of this emergent force by purely statistical arguments. This reflects the fact that the exact drift mechanism is irrelevant: Random drift or drift with a random component yields an entropic force. This lets newly formed engrams that are not in equilibrium undergo a transformation on the macroscopic scale. For the simple model examined in this section, the drift yields the only force behind this transformation. In the next sections we consider also the effects of neuronal connectivity preferences and structural connectivity, which lead to more complicated long-term dynamics than an exponential relaxation to equal coding levels.

### An engram energy

A hallmark of the interconnectivity of neurons in the brain is that it is generally sparser between neurons in different brain regions. This is a consequence of factors such as space and energy constraints, which limit the number of potential connections [39]. We will henceforth refer to two neurons as structurally connected if a synapse can potentially exist between them [40]. Whether a synapse exists between two structurally connected neurons is influenced by synaptic plasticity [41, 42], often in an activity dependent manner. To capture effectively the interplay between neuronal activity, synaptic plasticity, and structural connectivity, we again choose a bird’s eye, statistical perspective: We assign each microstate ***m*** an energy *H*(***m***). Microstates with lower energy are preferred by the engram and have a higher chance of occurrence. This allows us to describe the states of our neural system as a canonical ensemble [36], i.e. the probability of a microstate is given by the Boltzmann distribution, *p*(***m***) ∝*e*^−*βH*(***m***)^. This is in contrast to our first model, where all accessible microstates were equiprobable (have the same energy). In statistical physics, the constant *β* is the inverse of the temperature; small *β* (high temperature) indicates large random fluctuations and thus a strong entropic force. Analogously, in our system, *β* determines the strength of the randomness of the representational drift and thus the associated entropic force. In the absence of random representational drift (*β* →− ∞), the engram would always drift towards microstates with lower energy.

To construct an appropriate energy function, we introduce for a system consisting of *N* neurons an *N* × *N* structural connectivity matrix *A*, with element *A*_*ij*_ = 1 if neuron *j* can potentially form a synapse to neuron *i* and 0 otherwise. As before, the microstate ***m*** is a vector of length *N*, with component *m*_*i*_ = 1 if neuron *i* is an engram neuron and 0 otherwise. To write an expression for the energy we need to make assumptions about the nature of an engram; consistent with previous experimental and theoretical work, we assume that it is a neuronal assembly [43–45]: a group of strongly interconnected neurons. Each assembly neuron should have sufficient but not overwhelming recurrent input from the rest of the assembly. Therefore we introduce a constant *k* that represents an optimal, desired number of connections from other assembly neurons. Biologically, homeostatic plasticity and limited synaptic weights may increase the probability of such a configuration [31, 46] and thus implement the connectivity preference.

Furthermore, to foster an assembly, neurons should prefer to have reciprocal synapses. These are indeed more common than expected by chance in biological neural networks [47]; the preference could be implemented by symmetric forms of activity dependent plasticity [48]. We assume that all structurally permitted synapses between assembly neurons are formed. This is because we expect that biological assembly neurons tend to be coactive and that this leads to the strengthening of possible synapses. The above points lead us to assign the assembly-engram microstate the energy

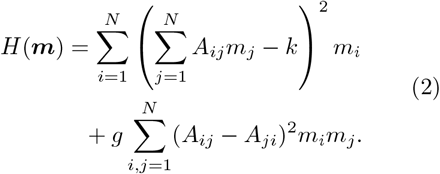

Its first term measures how much the number of synapses that each engram neuron receives from other engram neurons deviates from the desired number *k*. Such deviations are punished quadratically. However, this is not enough to favor a neuronal assembly: for example a circular feedforward chain [46, 49] would be likewise favored by this energy term. The distinguishing feature of an assembly are its reciprocal connections between the neurons. Missing reciprocal connections are thus punished by the second term. The constant *g* determines the relative strength of the two terms.

### Engram dynamics and (quasi-)equilibrium

To model the engram dynamics, we choose again in each time step randomly a neuron. If this neuron belongs to the engram, it may now stay or leave. The probability of either depends on which outcome is more favorable, i.e. on the resulting energy change. This can be interpreted as perturbing the state by preliminarily making the change and then probabilistically accepting or rejecting it. Outcomes more favorable than the current state, which result in a negative energy change, are more probable, Fig. 4a. Smaller *β* renders the energy change less relevant: it yields, for example, higher probabilities that the neuron leaves despite resulting less favorable states. This leads overall to stronger random drift. An alike probabilistic outcome selection happens if the neuron is originally not part of the assembly. Specifically we use the Glauber algorithm [50] for the simulation (Methods).

**Figure 4:**
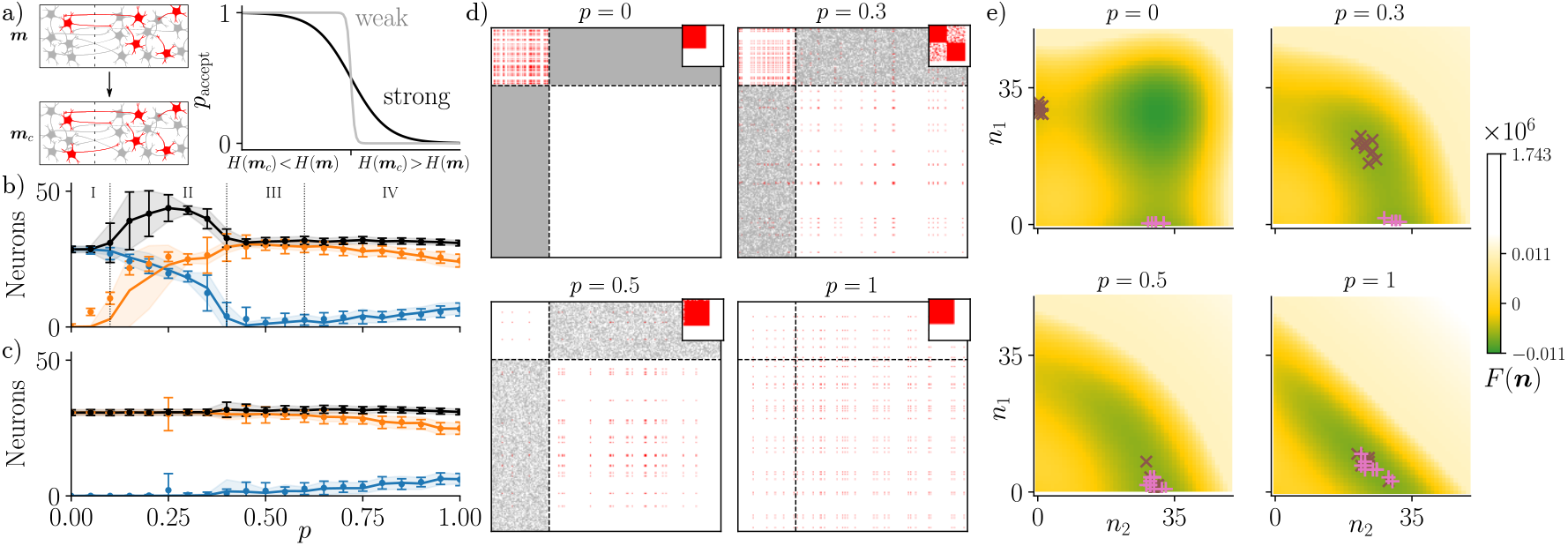
Engram dynamics and quasi-equilibria in the random-and-deterministic drift model (*β* = 0.012, *k* = 28, *g* = 5.5, *N*_1_ = 70, *N*_2_ = 280). (a) Energy-based simulation of engram drift. A candidate microstate is proposed by perturbing the current one, it is accepted or rejected depending on its energy. The probability of acceptance reflects the strength of representational drift. (b) Number of assembly neurons at quasi-equilibrium (mean ± std) in the first (blue) and the second (orange) region and total size (black) as a function of inter-region structural connectivity; initially the assembly is completely in Region 1. Points show mean ± std for the exact model, solid lines with shaded regions show the same for the simplified one. Dotted lines indicate parameter domains (I-IV) with different engram behavior. (c) Same as (b), but with the engram initially completely in Region 2. (d) An example of connections between assembly neurons after long time evolution, for four different levels of structural connectivity between the regions, as determined by the inter region-structural connection probability *p*. Initially the engrams are completely in Region 1. Connections between assembly neurons are shown in red, existing structural connections in white and absent structural connections in gray. Dashed lines indicate region boundaries. Reordering shows preserved engram structure (inset, focusing on non-empty matrix part) (e) Free energy *F* (***n***) as a function of the macrostate ***n*** for the same four levels of structural connectivity between the regions as in (d). Crosses indicate macrostates of an ensemble of engrams after long evolution (brown “×”: engrams initially completely in Region 1, pink “+”: engrams initially completely in Region 2). ***n*** of length *R* that assigns each region the number engram neurons in it. Starting with Eq. 2, the averaging yields for a microstate ***m*** that belongs to the macrostate ***n*** the energy (see Methods for details)

To illustrate the engram dynamics generated by our model, we first apply it to a network of two regions. The structural connectivity is specified by *p*_11_ = *p*_22_ = 1 and *p*_12_ = *p*_21_ = *p*, where *p*_*sr*_ is the probability that a structural connection from a neuron in region *r* to neuron in region *s* exists. Thus, within a single region each synaptic connection is possible and the probability of possible interregion connections is symmetric. We again begin with the engram initially located completely within one of the regions. Fig. 4b-d displays resulting very long-lived metastable states; transient dynamics for different *p* are shown in Fig. S1. The very long-lived metastable states can be quasi-equilibria, which would change for time tending to infinity, or true equilibria; if we do not need to emphasize this distinction, we denote them all as quasi-equilibria for simplicity. Remarkably, we observe that for the chosen energy function we can find parameters such that the statistical model has qualitatively the same behavior as the detailed biological model introduced in the next section.

If the engram is initially in the smaller region, Fig. 4b, there is an intricate dependence of its quasi-equilibrium on the regions’ interconnectivity *p*. It may be explained as follows: For *p* ≈ 0, parameter domain I, the regions are basically separated and the engram remains in the original region for the considered long simulation times. As *p* increases, domain II, neurons from the larger region can join the engram. The regions are, however, still only weakly interconnected and the synaptic contribution from one region to the other remains small. Therefore, each region contains on its own a slightly smaller number of engram neurons as a single region would host to maintain the desired level of interconnectivity. The slight reduction in the number of neurons is due to the input from the other region. As *p* increases further, domain III, the engram can freely drift between the two regions, and, since the engram energy is smaller the more reciprocal connections it has, it moves nearly completely to one region, choosing the entropically more favorable, larger one with more available microstates. As the probability *p* increases further, domain IV, the engram is again found in both regions, since the energy of such microstates is not prohibitively large anymore. As *p* tends to 1 there is less and less distinction between the regions and the system tends to behave like one large region; the quasi-equilibrium averages are thus determined by the entropy: the numbers of engram neurons are such that the coding levels in both regions become equal as in our first model. If the engram is initially in the larger region, the number of engram neurons in this region simply decays with increasing *p*. In particular, there is no analog to domain II, because there are too few neurons in region 1 to expect that some have by chance sufficiently many connections with assembly neurons of Region 2. Therefore practically no neurons in Region 1 join the assembly despite the considered long simulation times and it remains in Region 2.

### An engram free energy

Microstates that belong to the same macrostate can have different energies Eq. 2 due to the particular realization of the structural connectivity. In order to better understand the evolution of the assembly, we construct a simplified model by averaging the energy over the realizations of structural connectivity. For this we consider *R* brain regions and again denote the probability of a structural connection from a neuron in region *r* to a neuron in region *s* by *p*_*sr*_. A macrostate is then specified by a vector ***n*** of length *R* that assigns each region the number engram neurons in it. Starting with Eq. 2, the averaging yields for a microstate ***m*** that belongs to the macrostate ***n*** the energy (see Methods for details)

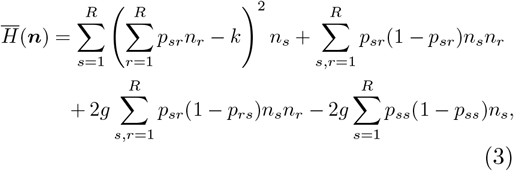

which only depends on the macrostate. Together with our assumption that the probability of a microstate is given by the Boltzmann distribution, this implies that the probability of macrostate ***n*** is proportional to its multiplicity Ω(***n***) times *e*^−*βH*(***n***)^, which is just the sum of the probabilities of the microstates associated to ***n***. We can rewrite this using the so-called free energy 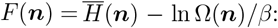 the probability of macrostate ***n*** is then proportional to *e*^−*βF* (***n***)^. The free energy accounts for the energy and the multiplicity of the macrostate. Thereby, in biological terms, it captures the interaction between neuronal activity and plasticity rules, represented by the energy function, on the one hand and the entropic force on the other hand. For the simplified, network realization-averaged model the (quasi-)equilibrium Fig. 4a,b as well as approach to it, Fig. S1, matches well that of the exact model. The simplified model explicitly highlights the contribution of the macrostate entropy term, *S*(***n***) = ln Ω(***n***), which exclusively governed the dynamics of our very first engram model, Fig. 3, and implicitly strongly impacts the engram evolution in the full energy model. In the presence of random representational drift, the engram tends to evolve towards minima of the free energy rather than the energy, Fig. 4c, like statistical physics systems that are described by a canonical ensemble [36]. For the chosen system with symmetric structural connectivity probability, the energy, Eq. 3, is symmetric under switching the number of neurons in regions. Thus, differences in reached quasi-equilibria between the regions, Fig. 4, are due to differences in the initial conditions and in the entropic force. The initial conditions in Fig. 4b vs. c and in Fig. 4e are mirrored between Regions 1 and 2 [(*n*_1_, *n*_2_) = (15, 0) vs. (*n*_1_, *n*_2_) = (0, 15)]. The symmetric energy function alone would thus induce mirrored temporal dynamics and mirrored quasi-equilibria: Fig. 4c would look like Fig. 4b with orange and blue colors interchanged and the sets of quasi-equilibria in Fig. 4e would be mirror symmetric along the diagonal. The entropic force breaks this mirror symmetry (due the the difference in region sizes), which manifests itself also in the strongly asymmetric free energy landscape, Fig. 4e. We note that depending on the inter-region connectivity *p* and the form of energy, the engram can be located in both regions or in only one. Since different engram types may have different forms of engram energy, this may explain why some engrams remain hippocampus-dependent while other do not.

### A biologically detailed model

In this section we develop a biologically detailed model of a single engram to examine its drift in a two-region network. As in the previous paragraph we model the engram as a neuronal assembly. This allows us to base our model on previous models of multiple, non-overlapping assemblies that drift in a single region [31, 32]: The network (see Methods) consists of linear Poisson (“Hawkes”) neurons, describing excitatory neurons in the balanced state. We use standard spike timing-dependent plasticity (STDP) with a slight modification: The strength of the STDP depends on the firing rate of pre- and postsynaptic neurons [51]. Further there is a rate dependent weight decay of input synapses [52].

We divide the network into two regions with full intra-region structural connectivity and symmetric inter-region connectivity probability, as in the previous section. Fig. 5a shows a resulting single drifting assembly in a network with two regions. Fig. 5b displays the average number of assembly neurons in the two regions after long evolution as a function of inter-region connectivity. Trajectories towards quasi-equilibrium are shown in Fig. S2. The qualitative features match the ones of the statistical model of the previous section.

**Figure 5:**
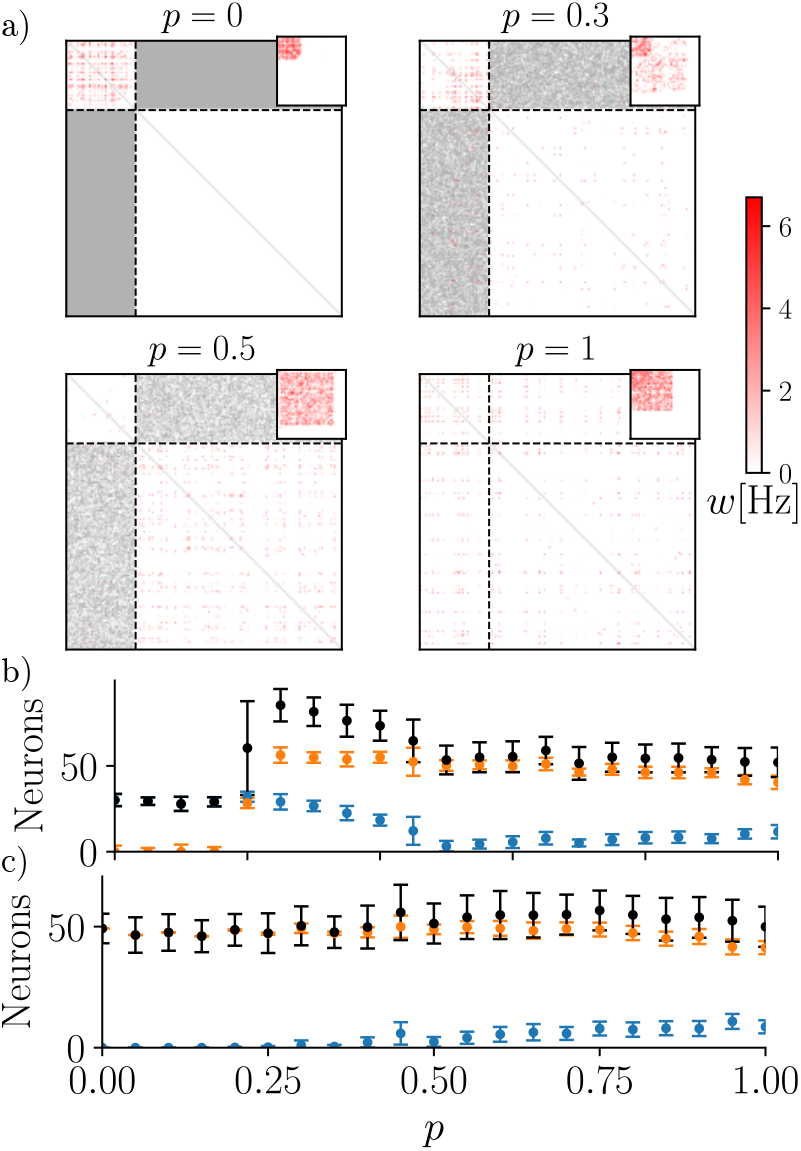
Quasi-equilibria in the biologically detailed engram drift model with two regions. (a) Example assembly weight matrices after long evolution for different inter-region connectivities, displayed as in Fig. 4b. (b,c) Number of assembly neurons (mean ± std) in Region 1 (blue) and 2 (orange) and total size (black), after long evolution. The assemblies are initially completely in (b) Region 1 or (c) Region 2. The dependence on interregion connectivity is similar as in the random-and-deterministic drift model, cf. Fig. 4b,c.

### Drifting engram in the mouse brain model

One advantage of our statistical model, especially the simplified, network realization-averaged version, is its relatively low computational cost. This allows to predict engram evolution in very large neural networks and even create a model of brain-wide engram drift. For this we combine brain-wide information about mouse neuron density and type [53], synaptic density [54], and connectivity [55,56], to construct a model of the mouse brain (Methods). Our model consists of 564 regions across two hemispheres. These regions are the mesoscale anatomical structures from the Allen Brain Atlas with sufficient available data. For each region we have estimated the number of excitatory neurons, *N*_*s*_, and the probability of these neurons to be structurally connected to excitatory neurons in other regions, *p*_*sr*_ (Methods). The engram evolution in this network is simulated like before: we use Glauber dynamics with the network realization-averaged energy function given by Eq. 3. As initial engram macrostate we take a fear memory engram distribution across the mouse brain that was experimentally measured three days after learning [10]. In the simplified model, Eq. 3, all microstates making up this macrostate are equiprobable; we randomly pick one of these microstates accordingly. Reached quasi-equilibria for different parameter values of the energy function are shown in Fig. S3. Some parameter combinations lead to “forgetting”: the engram completely vanishes. Others lead to very large coding levels for some regions. We consider such results as pathological and the parameters as invalid (Methods). Forgetting occurs especially for high numbers of desired inputs from other engram neurons, *k* = 10^4^ and *k* = 10^5^, as not enough such inputs are available.

Figure 6 shows the engram dynamics and their quasi-equilibria for some prominent brain regions for two valid parameter sets. Fig. S4 analogously displays results for the remaining valid parameter sets; Figure 6a,d,e shows coding levels in further brain regions. Figure 6 illustrates that the predictions for individual regions can differ: in one case we observe a substantial increase in engram neurons in the basolateral amygdala and in another a rapid increase in the anterior cingulate area. However, there are also typical, consistent predictions across valid parameter sets: For example, the engram quickly leaves the hippocampal fields CA1-3 that form large parts of the hippocampus. This fits classical ideas of memory consolidation (Introduction). Further the overlap between the current and the initial engram quickly decays to zero: the engram quickly drifts away completely. This indicates that some compensation that adjusts the inputs and outputs of the assemblies to conserve behavior must take place, such as unsupervised compensation [31]. Finally, the simulations predict that the overall engram is conserved in the sense that its size does not change substantially during its drift. The prediction of strongly different coding levels in different regions at quasi-equilibrium differs markedly from those obtained with our purely-random drift model and shows the relevance of connectivity and engram energy.

**Figure 6:**
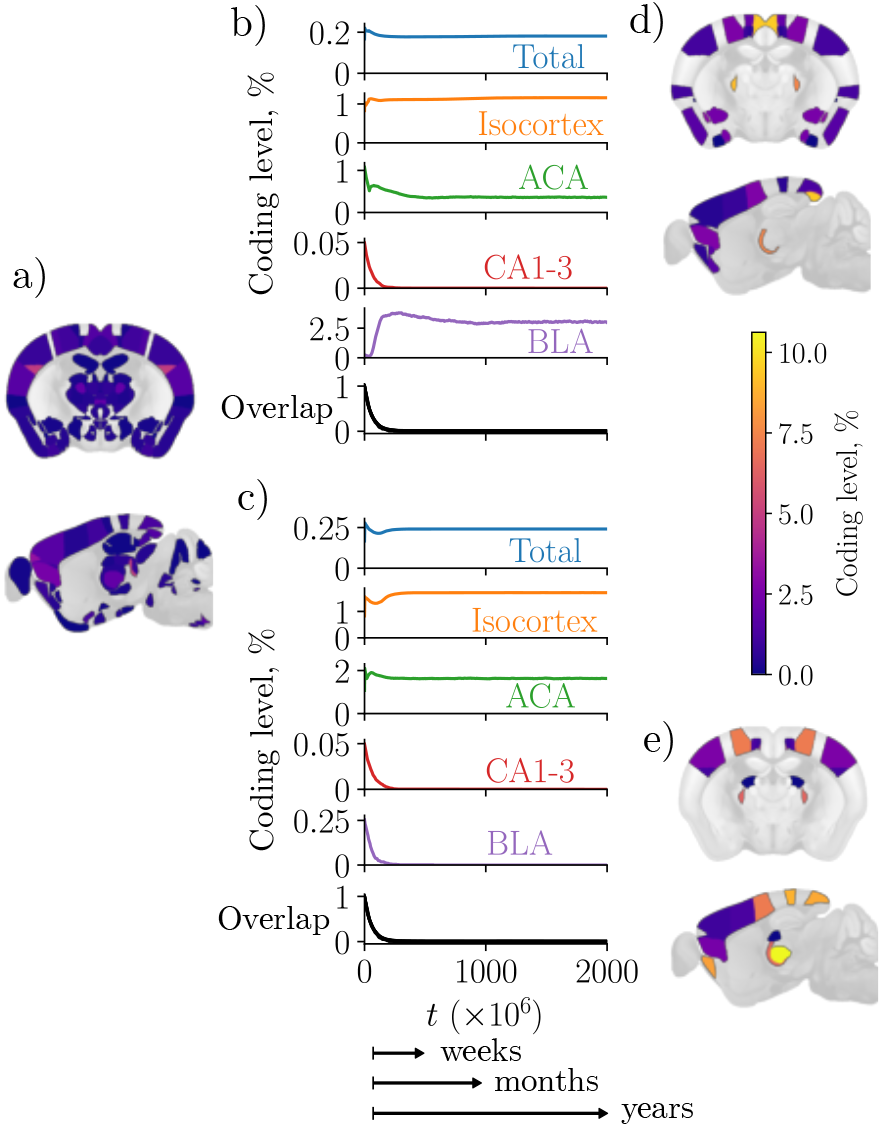
Fear memory engram dynamics and quasi-equilibria in the mouse brain. (a) Initial coding levels inferred from Ref. [10], coronal and sagittal sections, overlaid on Allen Mouse Brain Atlas [57] template. Coding levels for regions without engram neurons are not shown. (b) Total engram size, coding levels in selected macroscopic and mesoscopic regions and self-overlap dynamics for a valid parameter set (*β* = 0.01, *k* = 250, *g* = 0.1). Time *t* is Glauber steps; a suggested rough estimate of the timescale based on the decay of selfoverlap [3, 31] is displayed below. (c) Same as in (b) but for a different valid parameter set (*β* = 0.001, *k* = 500, *g* = 1). (d) and (e): Same as in (a) but for the quasi-equilibria of the parameter sets used in (b) and (c), respectively. BLA - basolateral amigdala; ACA - anterior cingulate area; CA1, CA2, CA3 - hippocampal fields.

## Discussion

We have studied how the presence of a memory engram in different regions of the brain dynamically changes due to representational drift that is purely random or deterministic with a random component. The results suggest that the process of memory consolidation may rely on such representational drift. The developed approach is general in following sense: Its fundamental implications, especially the emergence of an entropic force, apply whenever representational drift has a random component. The drift mechanism is thereby irrelevant. Further the transfer of an engram between regions does not rely on a specific network architecture. Finally the approach requires only minimal, generally accepted assumptions on the engram’s nature, namely that it is formed by neurons or by neurons and their interconnections.

The presence of randomness in representational drift is generally very likely because random fluctuations are ubiquitous in biological systems [58] and, in particular, in the nervous system [59]. Furthermore, several recent modeling studies [31,32,60–62] have shown that random representational drift can occur as a consequence of experimentally observed highly noisy spiking activity and random remodeling of synapses, in conjunction with activity dependent plasticity or homeostatic plasticity or both.

We have shown how to statistically describe the transformation of engrams over time. This accounts on the one hand for the entropic force that emerges due to random drift and the many possible engram states. On the other hand, there are forces that induce a deterministic drift. They may originate from plasticity mechanisms, which for example implement connectivity preferences; the (inhomogeneous) structural connectivity shapes them. To incorporate these forces, we introduced the concept of engram energy, which measures how beneficial an engram state is for the engram. The combination of random and deterministic drift is described by an engram free energy.

Our model allows to predict how engrams drift through the brain after learning. In general the entropic force drives the engram towards an equal distribution with identical coding levels across brain regions. This suggest that random representational drift may be employed by the brain to aid the creation of distributed memory representations. Furthermore, it allows engrams to go over potential barriers to more beneficial states. These functions add to previously suggested functions of the representational drift, which range from drift being a “bug” with no beneficial functions [63], to clearing space for memory storage [64, 65], regularization [66], sampling of solutions [67], and time stamping [68].

We derived our model from phenomenological considerations, for which the precise drift mechanisms are irrelevant. In particular, the origin of the random representational drift component is not important. Furthermore, detailed knowledge of plasticity mechanisms is not required. Rather, the allowed engram states and some preference characteristics are enough to construct the engram energy. A comparison with a biologically detailed model shows qualitative agreement between the predicted engram dynamics. Additional biological detail can be easily incorporated into our theory, like regional or temporal differences in neuronal excitability and in the magnitudes of random fluctuations.

Previous theoretical work on drifting assemblies studied drift within a single region [31–33]. Refs. [31, 32] model multiple drifting assemblies without overlaps, which completely tile the region. The drift is random and occurs due to noisy spiking activity, synaptic turnover and changes in spontaneous rate. Ref. [33] models a single assembly, which drifts due to a sequence of transient changes in excitability.

In our theory memory consolidation is a transformation of engrams from the initial non-equilibrium state towards stable (quasi-)equilibrium. This transformation is driven by both random and deterministic representational drift. Previous theoretical models of memory consolidation generally consider a few brain regions [17–26]. These models often use elaborate plasticity schemes to transfer memories from one region (usually a hippocampus model) to another (usually an isocortex model). In our model already a purely random drift can achieve such a transfer of memories from one region to another, if the size to the region to which engram is to be transferred is much larger. The deterministic drift (determined by the engram energy) modifies the transfer dynamics and resulting engram states, but preserves the general description of memory consolidation as drift.

Our study proposes an entirely novel, statistical physics-based class of models for representational dynamics. These models are both analytically and computationally well tractable. The latter allowed us to simulate the brain-wide engram transformation of fear memory in the mouse brain. This resulted in detailed experimental predictions. There is some uncertainty in the predictions resulting from uncertainty about the parameters of our energy function, which will be eliminated when the predictions at a single or a few time points are experimentally tested. Our models further enable future *in silico* experiments that predict the effects of perturbations to the engram or the neural network. For example, it is possible to systematically lesion different brain regions and examine resulting engram dynamics.

We do not explicitly take into account the contributions of inhibitory neurons to the engram. However, our approach could be extended by incorporating terms into the energy function that include inhibitory engram neurons.

We expect our approach to be applicable to various specific types of memories (e.g. episodic, semantic, motor). In particular, we expect that it will be possible to construct energy functions for specific types of engrams, for example sequentially structured ones, to predict their long term drift. Furthermore, it will be possible to add terms to the energy that account for the interaction between different engrams.

## Methods

### Purely-random drift model

To simulate engram evolution in our first model, Figs. 1-3, we replace in each simulation step a random engram neuron with a random non-engram neuron, in the sense that the picked engram neuron becomes a non-engram neuron and the picked non-engram neuron becomes an engram neuron. Each engram neuron has a *n/N* chance of being picked for replacement, and each non-engram has a (*N* − *n*)*/N* chance of replacing it.

We can write the dynamics of the macrostate of the two-region model in the form of a Markov chain,

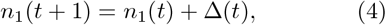

where *n*_1_(*t*) is the number of neurons in Region 1 at simulation step *t*. The increment Δ(*t*) can assume the values 0, 1, and − 1: If an engram neuron is replaced by a non-engram neuron from the same region, we have Δ(*t*) = 0, since the number of engram neurons in Region 1 does not change. If an engram neuron from Region 2 is replaced with a non-engram neuron from Region 1, the number of engram neurons in Region 1 increases by 1 and we have Δ(*t*) = 1. If an engram neuron from Region 1 is replaced with a non-engram neuron from Region 2, Δ(*t*) = − 1. The increments have the following probabilities of occurrence:

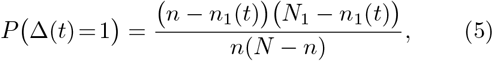

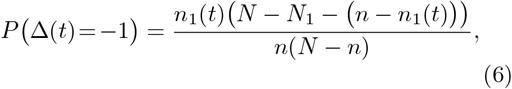

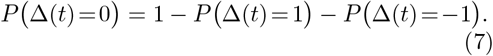

This is because *n*(*N − n*) is the total number of ways that we can combine (and thus replace) one of the *n* engram neurons with one of the *N* − *n* non-engram neurons; further, for example (*n* ™ *n*_1_(*t*))(*N*_1_ ™ *n*_1_(*t*)) is the number of ways in which we can combine (replace) one of the *n* − *n*_1_(*t*) engram neurons of Region 2 with one of the *N*_1_ ™ *n*_1_(*t*) non-engram neurons of Region 1. We note that these considerations straightforwardly generalize to multi-region models. In fact, if we focus on the macrostate of region 1 as above in such a model, the same formulas hold as above, because we can gather all other regions into one region.

The Markov chain specified by Eqs. 4 to 7 is irreducible and aperiodic and thus has a unique equilibrium distribution [36]. We can rewrite the macrostate equilibrium probability *p*(***n***) = Ω(***n***)*/*Ω(*n*) as function of the single variable 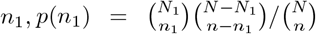, using the fact *n*_2_ = *n* – *n*_*1*_ We obtain 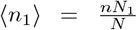 and 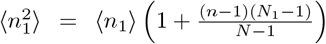 by applying the definition of expectation directly and using the Chu–Vandermonde identity. For *n*_1_, *N*_1_ ≫ 1, 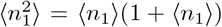, such that the coefficient of variation is ⟨*n*_1_⟩ ^−1*/*2^.

To derive the analytical expression for the average trajectory Eq. 1, we calculate the expected number of engram neurons in Region 1 after one simulation step; in other words, we compute the conditional expectation of *n*_1_(*t*) given *n*_1_(*t* − 1),

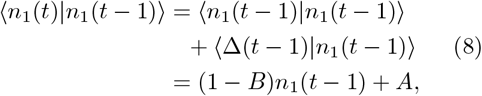

where 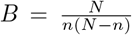 and 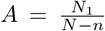, and we have used Equations (5) to (7) to directly average the second term. We obtain for *u* > *t*

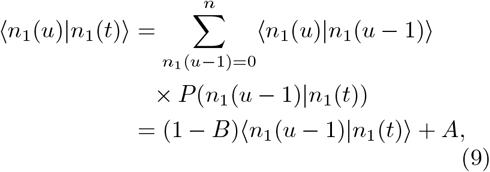

where we used the Markov property, which implies that *n*_1_(*u*) depends on *n*_1_(*t*) only via *n*_1_(*u* ™ 1), and Eq. 8; the detailed derivation is given in Supplemental section S1. Starting with some initial state *n*_1_(0) and applying *t* times the recurrence relation given by Eq. 9 yields Eq. 1.

### Random-and-deterministic drift model

At each simulation step we construct a candidate microstate, ***m***_***c***_, by modifying the current microstate ***m*** as follows: We pick a random neuron. If this neuron is an engram neuron, we remove it from the engram; if the picked neuron is a non-engram neuron, we add it to the engram. The candidate microstate is accepted with probability 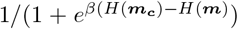, otherwise, with probability 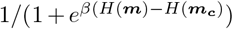, the system stays in the current microstate. An analogous procedure is applied for the simplified model; in this model the energies of different microstates are equal if they belong to the same macrostate.

To obtain Eq. 3 we average Eq. 2 over the realizations of the *A*_*ij*_

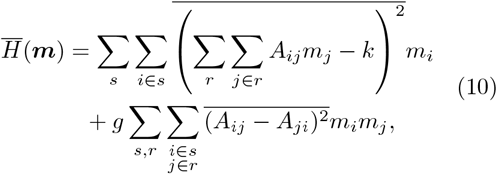

where indices *i* and *j* run over neurons and *s* and *r* over regions, *k* ∈ *q* signifies that neuron *k* is in region *q*. To convert from microstates to macrostates we use 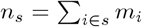. We allow autapses, i.e. it is possible that *A*_*ii*_ = 1. Expanding terms of Eq. 10, averaging using the fact that *A*_*ij*_ are independent Bernoulli random variables, and rearranging leads to Eq. 3; the detailed derivation is given in Supplemental section S2.

### Biologically detailed assembly model

The network consists of *N* linear Poisson neurons that spike with instantaneous rates *f*_*i*_(*t*), for *i* = 1, 2, …, *N* . The spike rate of a linear Poisson neuron is incremented instantaneously at the arrival of an input spike by an amount proportional to the synaptic weight. In the absence of input spikes the rate decays exponentially with time constant *τ* to the background value *f*_sp_. The dynamics are thus given by

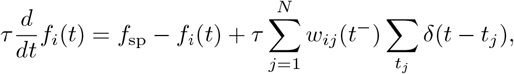

where *w*_*ij*_(*t*^−^) is the strength of the synapse from neuron *j* to neuron *i* just before time *t* (where it may change in jump-like manner, see below); *t*_*j*_ are the spike times of neuron *j*. The synaptic weights are positive, with maximum possible weight *w*_max_: 0 *< w*_*ij*_ *< w*_max_.

The synaptic strengths are modified at each pre- and each postsynaptic spike by STDP that depend also on the rates of pre- and post synaptic neurons

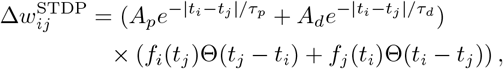

where *A*_*p*_, *A*_*d*_, *τ*_*p*_, *τ*_*d*_ and Θ is the Heaviside step function. In addition, each synaptic weight decays at a constant rate. Finally if a neuron *j* spikes, its output synaptic weights are instantaneously decremented by an amount proportional to its frequency,

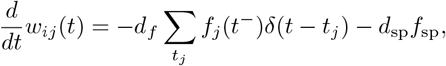

where *d*_*f*_ and *d*_sp_ are constants, and the sum is over all spikes times *t*_*j*_ of neuron *j*.

### Mouse brain model

The regions in our model are “summary structures” - non-overlapping mesoscale brain regions defined by the Allen Mouse Brain Atlas [57]. We use subscripts *r* and *s*, ranging from 1 to *R*, to index these mesoscopic regions. The Allen Mouse Brain Atlas also defines 12 macroscopic brain regions [57]: isocortex, olfactory areas, hippocampal formation, cortical subplate, striatum, pallidum, thalamus, hypothalamus, midbrain, pons, medulla, and cerebellum. We use this macroscopic structuring to obtain reference regions for the estimation of unknown data (synaptic densities) and to assign long range connectivity properties (see below).

We obtain each mesoscopic region’s volume *V*_*r*_ and number of excitatory *N*_*r*_ and inhibitory 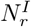 neurons from Ref. [53]. The structural connectivity in the model is based on the mesoscale connectome obtained by Ref. [56] and refined by Ref. [55]: These studies measured connection strengths by anterograde tracing such that both excitatory and inhibitory projections are accounted for. We obtain the volume-normalized connection strength between two regions, 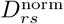, from ref. [55] and convert it to the connection strength 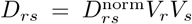. We are interested in the excitatory connection strengths, 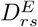, only. We assume that all long-range (inter-region) connections are excitatory, 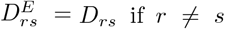, with the exception of those that are part of the subcortical brain regions and cerebellum, which have prominent inhibitory projections [69]. We therefore assume that the number of excitatory inter-region projections is proportional to the number of excitatory neurons 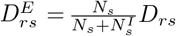, if the region *s* belongs to the subcortical regions, i.e. to the macroscopic regions cortical subplate, striatum, pallidum, thalamus, hypothalamus, midbrain, pons, medulla, or if the region belongs to the cerebellum excluding the cerebellar cortex. The cerebellar cortex has only inhibitory outputs [70], and we therefore set 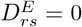 if *s* is a region in the cerebellar cortex. For the intra-region connections, we estimate the connection strength due to excitatory projections 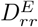 by assuming that the fraction of excitatory intra-region connections equals the fraction of excitatory neurons, which yields 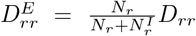. We assume that the number of excitatory synapses from region *s* to region *r, S*_*rs*_, is proportional to the strength of connections as measured by the anterograde tracing 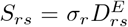. We estimate the region-specific proportionality constant *σ*_*r*_, in a method similar to the one used in Ref. [71]: Summing the number of excitatory synapses from all input regions *s* to a region *r*, must yield the total number of excitatory synapses in that region,

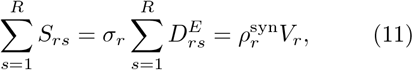

Where 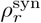, is the density of excitatory synapses in region *r*, which we obtain for some regions from Ref. [54]. For regions for which it is not available, we estimate 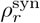 by taking the average over regions in the same macroscopic region with known densities. Solving Eq. 11 for *σ*_*r*_ and inserting the result into the defining equation for *S*_*rs*_ yields

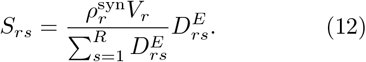

Assuming that all 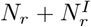 neurons in a region *r* are statistically equivalent, the probability that a particular synaptic input from a region *s* is present is given by the number of synaptic inputs from region *s, S*_*rs*_, divided by the number 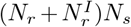 of in principle possible connections. For our simulations we consider only the excitatory neurons. The probability 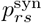 that an excitatory neuron in region *r* receives a synapse from an excitatory neuron in region *s*, is then also given by

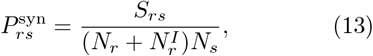

for region *r* with excitatory neurons (*N*_*r*_ > 0), otherwise 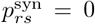 . Here we accounted for the fact that we only considered output connections of excitatory neurons, but did not distinguish whether they end at excitatory or inhibitory neurons. The total number of possible connections that we divide by is therefore the product of the number of excitatory neurons in the sending and the total number of neurons in the receiving area.

The probability 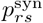 calculated above relates to the already existing synapses. We assume that the probability of a structural (potential) connection is proportional to it, 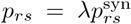. The proportionality constant *λ* depends on the filling fraction and the number of synapses formed between two neurons. The filling fraction estimates the ratio of existing synapses to all potential ones without major axonal or dendritic remodeling. It is estimated to be 0.26 in some areas of mouse isocortex, and ranges between 0.12 and 0.34 for different brain areas and model organisms [40]. There is typically more than one synapse between connected neurons. For example, for the rat isocortex on average 5.5 [72] and 4.7 [73] synaptic contacts between neurons were reported. Multiple synapses were also reported for the pairs of neurons form the mouse isocortex [74]. Taking into account multiple synapses between pairs of neurons on average compensates the effect of the filling fraction, and we therefore set *λ* = 1.

We interpret *p*_*rs*_ as the probability that a synapse may in principle exist between two neurons, i.e. that these neurons are structurally connected. *p*_*rs*_ and *N*_*r*_ specify our brain model; it covers 282 brain regions (per hemisphere).

We obtain the initial state of the engram in region *s* as

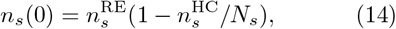

where 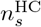 and 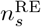 are the average numbers of excitatory neurons that were labeled by cFos in the home cage and during fear memory recall three days after fear conditioning, respectively [10]. The rationale behind this formula is as follows: The probability that a neuron is active in the home cage is 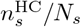 and the probability that a neuron is active during recall is 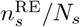. We expect that there is a background of spuriously active neurons that we need to subtract from 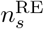 to get the true engram size. Due to the lack of information about these neurons and the expectation that spurious activity may be similar in both the recall and the homecage measurement, we use as a rough estimate that the number of spuriously active neurons equals the number of neurons that lie in the overlap of the sets of neurons that are active during recall and in the homecage. Assuming (somewhat contradictorily) that the probability that a neuron is active during recall and in the home cage are independent because the two events have little in common, the probability of a neuron to be spuriously active during a measurement is then 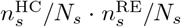. This yields as the probability of a neuron to be a true recall engram neuron 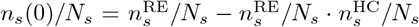 and thus for the expected number of recall engram neurons Eq. 14. We assume that both excitatory and inhibitory neurons are labeled by cFos with the same probability, and scale the experimental data by the fraction of excitatory neurons in the region to obtain only the excitatory engram.

We repeated simulations five times for each parameter set (*β, k, g*). We consider a set of parameters as pathological, if it led to a coding level of more than 20% in a region with more than 1000 excitatory neurons, at any simulation step, for the majority of realizations.

## Acknowledgements

We thank Simon Altrogge for enlightening discussions and insightful ideas, the German Federal Ministry of Education and Research (BMBF) for support via the Bernstein Network (Bernstein Award 2014, 01GQ1710), and the German Research Society (DFG) for support via a Research Grant (430156448).

