## Supplemental material for "Brain-wide representational drift: memory consolidation and entropic force"

### Supplementary figures

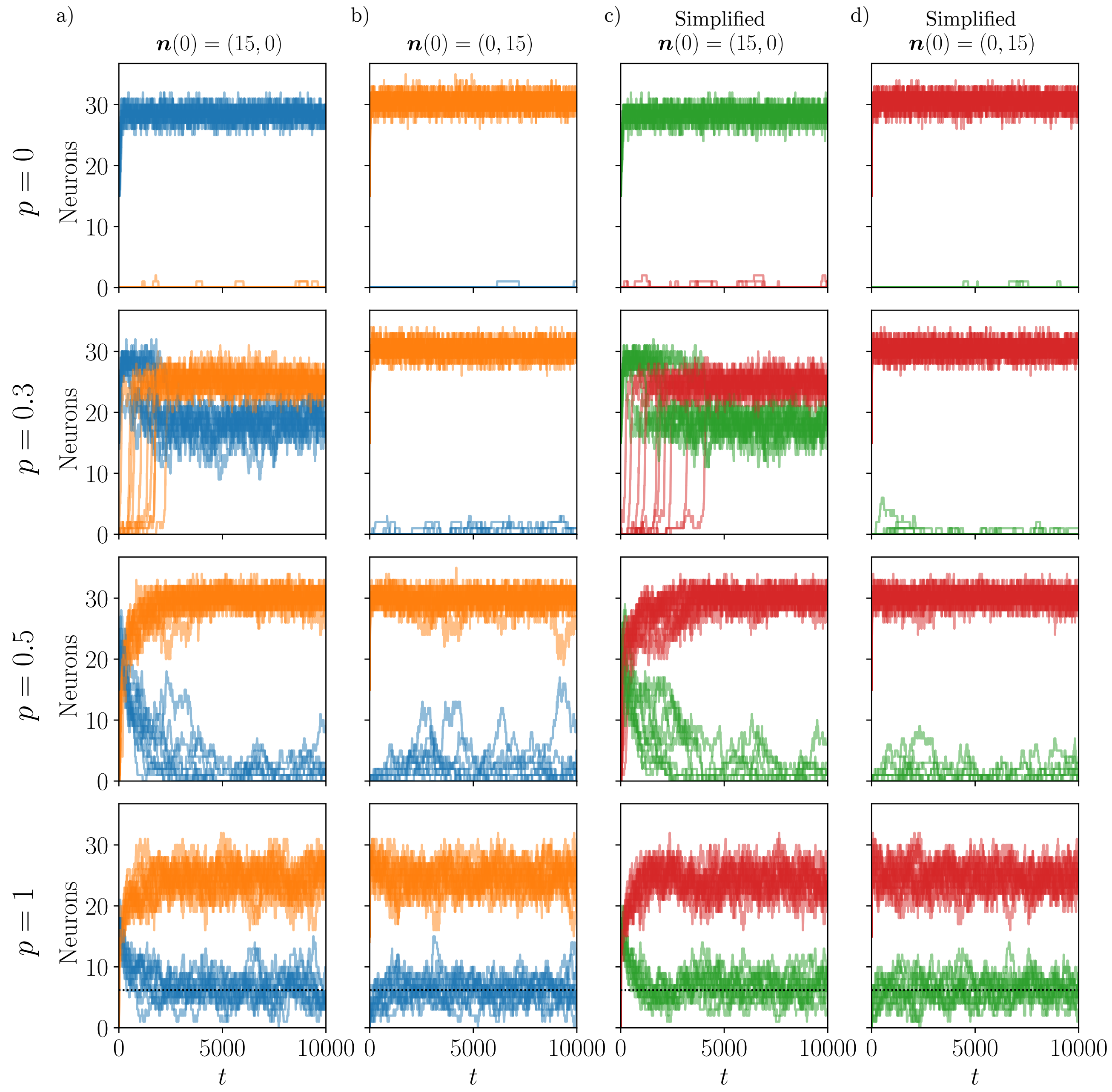


Figure S1: Engram macrostate dynamics in the random-and-deterministic drift model with full and simplified energy function (parameters as in main text Fig. 4). Left hand side columns a,b: dynamics in the model with engram energy given by main text Eq. 2. Right hand side columns c,d: dynamics in the simplified model with engram energy given by main text Eq. 3. Columns a,c: engram initially in Region 1; columns b,d: engram initially in Region 2. Rows: various inter-region connectivity probabilities. The number of neurons in Region 1 is displayed in blue (full model) or green (simplified model), the number of neurons in Region 2 in orange (full model) or red (simplified model). Ten realizations are shown for each initial condition and probability. The dynamics in both models are qualitatively similar.

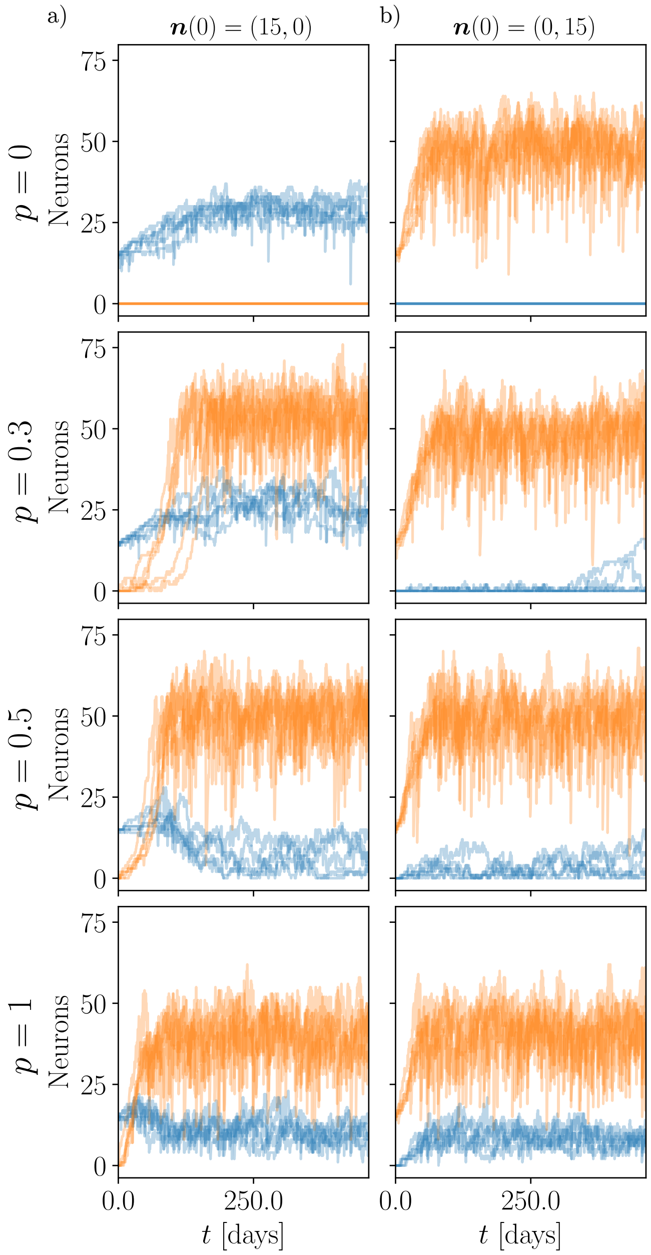


Figure S2: Engram macrostate dynamics in the biologically detailed model. Column a: engram initially in Region 1; columns b,d: engram initially in Region 2. Rows: various inter-region connectivity probabilities. The number of neurons in Region 1 is displayed in blue, the number of neurons in Region 2 in orange. We observe qualitatively similar forms of dynamics as in the random-and-deterministic drift model (cf. Fig. S1). There are also obvious differences: in particular the noise level in the biologically more detailed model is generally higher.

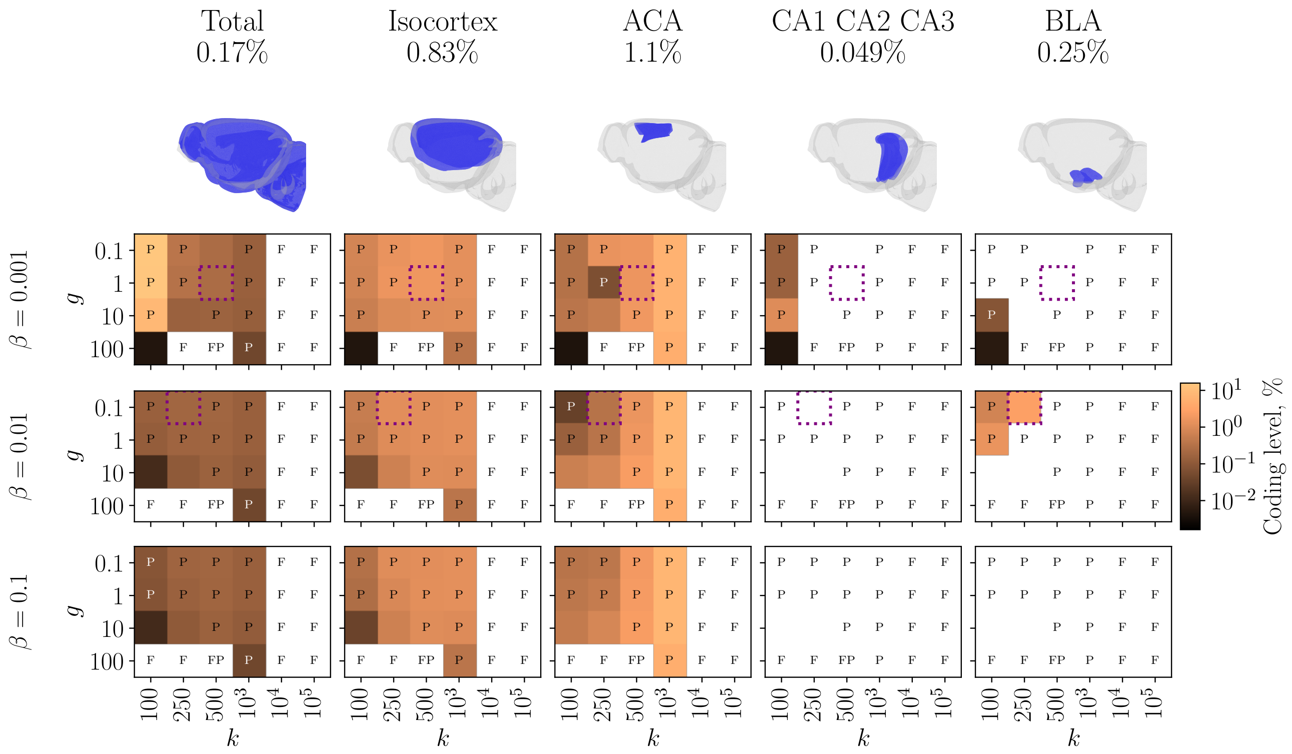


Figure S3: Quasi-equilibria of the fear memory engram in the mouse brain for different parameters of the energy. Columns: selected regions. Top row: schematic display of the region along with its initial coding level. Subsequent rows: Various parameters $\beta$. Matrices: Various parameters $g$ and $k$. Matrix entries: quasi-equilibrium coding levels, averaged over realizations. Zero coding level is shown in white. Purple-dashed squares indicate parameters shown in the main text Fig. 6. F - forgetting; P - pathological evolution; BLA - basolateral amigdala; ACA - anterior cingulate area; CA1, CA2, CA3 - hippocampal fields.

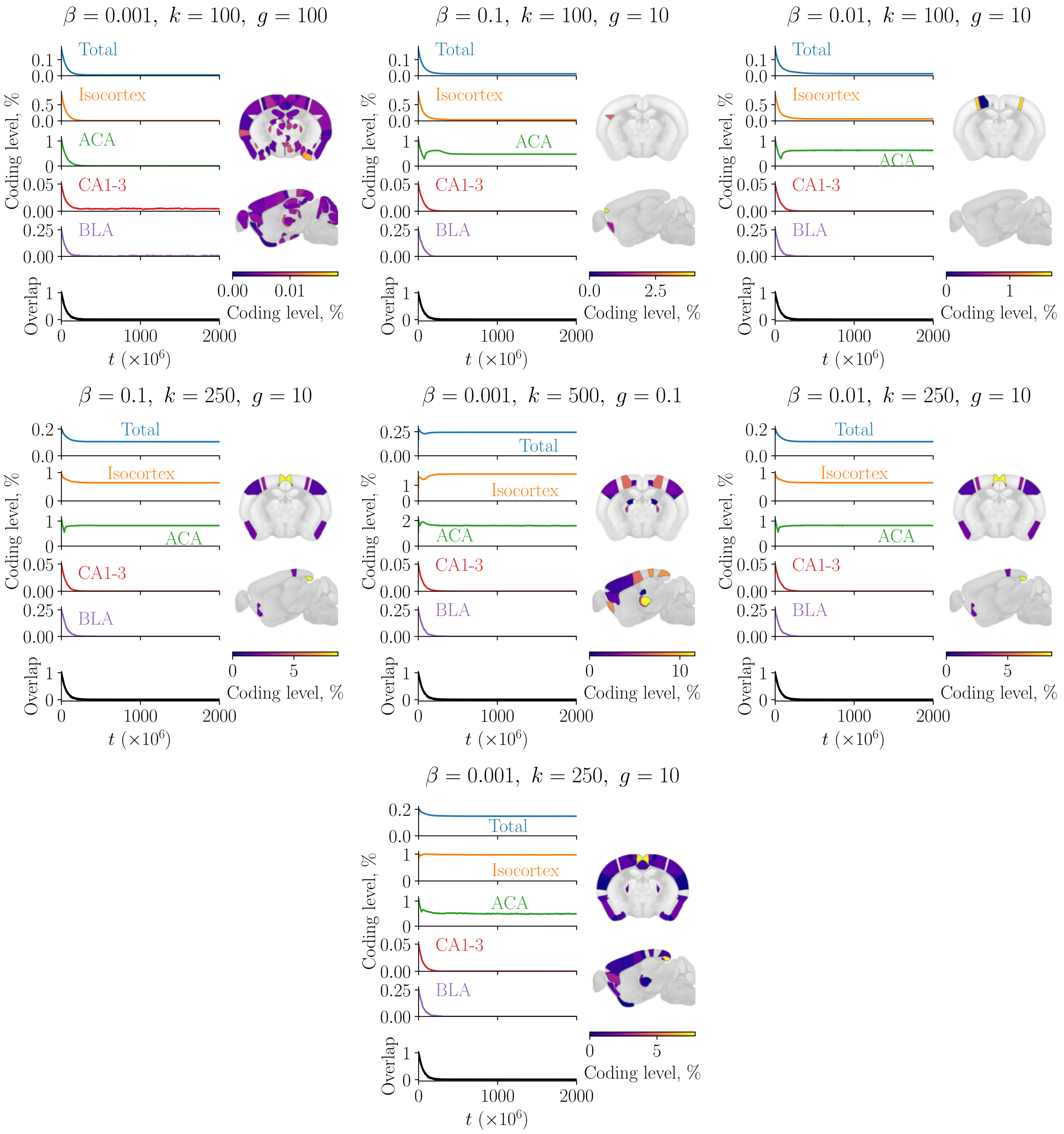


Figure S4: Fear memory engram dynamics and quasi-equilibria in the mouse brain as in main text Fig. 6, for the remaining valid parameter sets.

### 1 Derivation of the average trajectory

Main text Eq. 9 may be derived as follows:

$$\begin{aligned}
\langle n_1(u)|n_1(t)\rangle &= \sum_{n_1(u)=0}^{n} n_1(u)P(n_1(u)|n_1(t)) \\
&= \sum_{n_1(u)=0}^{n} n_1(u) \sum_{n_1(u-1)=0}^{n} P(n_1(u)|n_1(u-1))P(n_1(u-1)|n_1(t)) \\
&= \sum_{n_1(u-1)=0}^{n} \langle n_1(u)|n_1(u-1)\rangle P(n_1(u-1)|n_1(t)) \\
&= \sum_{n_1(u-1)=0}^{n} \big((1-B)n_1(u-1)+A\big)P(n_1(u-1)|n_1(t)) \\
&= (1-B)\langle n_1(u-1)|n_1(t)\rangle + A,
\end{aligned} \tag{1}$$

### 2 Derivation of the average energy

In the following we average main text Eq. 10 term by term. Since $A_{ij} \in \{0,1\}$ are independent Bernoulli random variables, we have $\overline{A_{ij}} = \overline{A_{ij}^2} = p_{ij}$. Furthermore, $\overline{A_{ij}A_{gh}} = p_{ij}p_{gh}$ if either $i \neq g$ or $j \neq h$ or both. We assume that all neurons in region $s$ have the same probability of being structurally connected to neurons in region $r$, that is $p_{ij} = p_{sr}$ for neuron $i$ in region $s$ and neuron $j$ in region $r$. We average the first term as follows

$$\begin{aligned}
\overline{\sum_s \sum_{i\in s}\left(\sum_r \sum_{j\in r} A_{ij}m_j - k\right)^2 m_i} &= \sum_{s,r,q} \sum_{\substack{i\in s\\ j\in r\\ h\in q}} \overline{A_{ij}A_{ih}} m_i m_j m_h - 2k \sum_{s,r} \sum_{\substack{i\in s\\ j\in r}} \overline{A_{ij}} m_i m_j \\
&\quad + k^2 \sum_s \sum_{i\in s} m_i \\
&= \sum_{\substack{s,r,q\\ r\neq q}} \sum_{\substack{i\in s\\ j\in r\\ h\in q}} p_{ij}p_{ih} m_i m_j m_h + \sum_{s,r} \sum_{\substack{i\in s\\ j,h\in r\\ j\neq h}} p_{ij}p_{ih} m_i m_j m_h \\
&\quad + \sum_{s,r} \sum_{\substack{i\in s\\ j\in r}} p_{ij} m_i m_j^2 - 2k \sum_{s,r} \sum_{\substack{i\in s\\ j\in r}} p_{ij} m_i m_j \\
&\quad + k^2 \sum_s \sum_{i\in s} m_i \\
&= \sum_{\substack{s,r,q\\ r\neq q}} p_{sr}p_{sq} n_s n_r n_q + \sum_{s,r} p_{sr}^2 n_s n_r (n_r - 1) \\
&\quad + \sum_{s,r} p_{sr} n_s n_r - 2k \sum_{s,r} p_{sr} n_s n_r + k^2 \sum_s n_s \\
&= \sum_s \left(\sum_r p_{sr} n_r - k\right)^2 n_s + \sum_{s,r} p_{sr}(1-p_{sr}) n_s n_r.
\end{aligned} \tag{2}$$

For averaging the second term note that it is zero if $i = j$, so we need to consider only the case $i \neq j$

$$\begin{aligned}
g\sum_{s,r}\sum_{\substack{i\in s\\ j\in r}}\overline{(A_{ij}-A_{ji})^2}m_im_j &= g\sum_{s,r}\sum_{\substack{i\in s\\ j\in r}}\left(\overline{A_{ij}^2}-2\overline{A_{ij}A_{ji}}+\overline{A_{ji}^2}\right)m_im_j \\
&= g\sum_{s,r}\sum_{\substack{i\in s\\ j\in r\\ i\neq j}}(p_{ij}-2p_{ij}p_{ji}+p_{ji})m_im_j \\
&= g\sum_{s,r}(p_{sr}-2p_{sr}p_{rs}+p_{rs})\sum_{\substack{i\in s\\ j\in r\\ i\neq j}}m_im_j \\
&= g\sum_{\substack{s,r\\ s\neq r}}(p_{sr}-2p_{sr}p_{rs}+p_{rs})\sum_{\substack{i\in s\\ j\in r}}m_im_j \\
&\quad + g\sum_{s}(2p_{ss}-2p_{ss}^2)\sum_{\substack{i,j\in s\\ i\neq j}}m_im_j \\
&= g\sum_{\substack{s,r\\ s\neq r}}(p_{sr}-2p_{sr}p_{rs}+p_{rs})n_sn_r \\
&\quad + g\sum_{s}(2p_{ss}-2p_{ss}^2)n_s(n_s-1) \\
&= g\sum_{s,r}(p_{sr}-2p_{sr}p_{rs}+p_{rs})n_sn_r - 2g\sum_{s}p_{ss}(1-p_{ss})n_s.
\end{aligned} \tag{3}$$

Combining the two terms yields main text Eq. 3.
